# Computational metabolomics illuminates the lineage-specific diversification of resin glycoside acylsugars in the morning glory (Convolvulaceae) family

**DOI:** 10.1101/2021.08.20.457031

**Authors:** Lars H. Kruse, Alexandra A. Bennett, Elizabeth H. Mahood, Elena Lazarus, Se Jin Park, Frank Schroeder, Gaurav D. Moghe

**Author notes:** Both authors contributed equally to the study (Lars H. Kruse, Alexandra A. Bennett).

## Abstract

Acylsugars are a class of plant defense compounds produced across many distantly related families. Members of the horticulturally important morning glory (Convolvulaceae) family produce a diverse sub-class of acylsugars called resin glycosides (RGs), which comprise oligosaccharide cores, hydroxyacyl chain(s), and decorating aliphatic and aromatic acyl chains. While many RG structures are characterized, the extent of structural diversity of this class in different genera and species is not known. In this study, we asked whether there has been lineage-specific diversification of RG structures in different Convolvulaceae species that may suggest diversification of the underlying biosynthetic pathways. Liquid chromatography coupled with tandem mass spectrometry (LC-MS/MS) was performed from root and leaf extracts of 26 species sampled in a phylogeny-guided manner. LC-MS/MS revealed thousands of peaks with signature RG fragmentation patterns with one species producing over 300 signals, mirroring the diversity in Solanaceae-type acylsugars. A novel RG from *Dichondra argentea* was characterized using Nuclear Magnetic Resonance spectroscopy, supporting previous observations of RGs with open hydroxyacyl chains instead of closed macrolactone ring structures. Substantial lineage-specific differentiation in utilization of sugars, hydroxyacyl chains, and decorating acyl chains was discovered, especially among *Ipomoea* and *Convolvulus* – the two largest genera in Convolvulaceae. Adopting a computational, knowledge-based strategy, we further developed a high-recall workflow that successfully explained ~72% of the MS/MS fragments, predicted the structural components of 11/13 previously characterized RGs, and partially annotated ~45% of the RGs. Overall, this study improves our understanding of phytochemical diversity and lays a foundation for characterizing the evolutionary mechanisms underlying RG diversification.

## Introduction

The plant kingdom has historically been a source of numerous metabolites of use in medicines, foods, cosmetics, agriculture, and cultural practices. Many of these metabolite classes e.g., glucosinolates, alkaloids, terpenes, acylsugars, or cardenolides show lineage-specific structural diversification. Such specialized metabolites play important functional roles in plant pollination, defense, and resilience to abiotic stress conditions in the respective plants’ ecological niche. One such class of specialized metabolites is acylsugars, which, in the Solanaceae family, are produced in the trichomes of many species (Fan et al., 2019; Landis et al., 2021; Leong et al., 2020, 2019; Moghe et al., 2017). In Solanaceae, this class of anti-herbivory compounds comprises a sugar core (usually a sucrose but sometimes glucose or inositol) to which acyl chains derived from fatty acid or branched amino acid metabolism are esterified **(Fig. 1A)** (Leong et al., 2020, 2019; Moghe et al., 2017). A previous study catalogued acylsugar diversity across 40 species in the family, and identified some species, such as *Salpiglossis sinuata* and *Nicotiana alata*, that produce over 600 acylsugars on individual leaves (Moghe et al., 2017). Acylsugars also show variation among individuals of the same species (Landis et al., 2021). This extreme structural diversity of acylsugars is hypothesized to occur due to the promiscuity of the BAHD acyltransferases involved in their biosynthesis, and the diverse precursor coenzyme A-activated acyl groups in the trichome metabolite pool.

**Figure 1:**
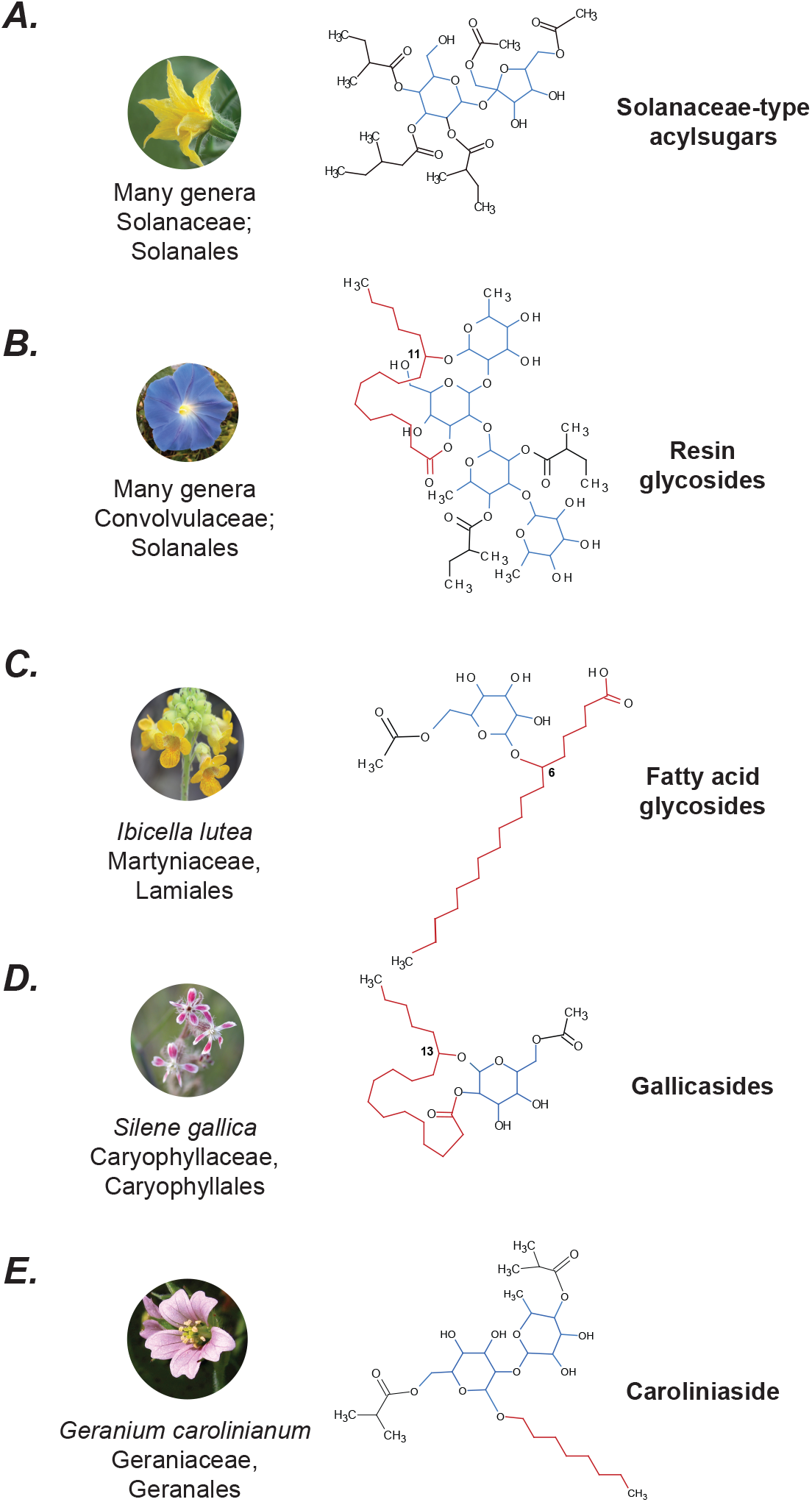
Representative structures of acylsugars from various species groups. referred to by the names in associated publications (see Main Text). The blue, red and black substructures refer to the sugar cores, hydroxyacyl chains and smaller ester decorations, respectively. Resin glycosides show the most complex structural patterns among the different known types of acylsugars shown here. Plant images under Creative Commons licenses from Wikimedia Commons.

While “acylsugars” commonly refers to the class of compounds that is known to be restricted only to members of the Solanaceae family, other structurally and possibly biosynthetically analogous acylated sugar compounds are also documented across the plant kingdom. For example, *Ibicella lutea* (Martyniaceae) (Asai et al., 2010), *Silene gallica* (Caryophyllaceae) (Asai and Fujimoto, 2010), *Cerastium glomeratum* (Caryophyllaceae) (Asai et al., 2012) and *Geranium carolinianum* (Geraniaceae) (Asai et al., 2011) are also known to produce acylated sugar compounds **(Fig. 1C–E)**. In the known examples, the central sugar in these compounds is a glucose, with acyl or hydroxyacyl chains attached via ester and/or ether linkages. Given all these compound classes are technically acylsugars, we refer to Solanaceae acylsugars as “Solanaceae-type acylsugars” while referring to the combined class as “acylsugars” **(Fig. 1)**. While the extent of the diversity of acylsugars across plants is not known, structures of another similar compound class – resin glycosides (RGs) – have been extensively characterized in morning glories (Convolvulaceae) since their discovery in the 1990s (Anaya et al., 1990; Pereda-Miranda et al., 1993) **(Fig. 1B)**. Over 300 RG structures have been characterized from several Convolvulaceae species using Nuclear Magnetic Resonance (NMR) (reviewed in (Eich, 2008; Pereda-Miranda et al., 2010)), and they reveal three primary structural elements: (i) an oligosaccharide core of 2-7 sugars comprising pentoses, deoxyhexoses and hexoses, (ii) a 14-18 carbon long hydroxy- or dihydroxy fatty acid that may form an intra-molecular macrolactone ring, and (iii) decorations of aromatic or aliphatic acyl side chains that can be 2-16 carbons long. Some RGs undergo intermolecular condensation to form larger, more complex units (Eich, 2008; Pereda-Miranda et al., 2010). RGs have also been extensively characterized functionally, with studies finding their roles in bacterial and fungal inhibition (Harrison et al., 2003), in herbivory defense against nematodes (Harrison et al., 2003; Jackson and Peterson, 2000) and leaf herbivores (Jackson and Peterson, 2000) as well as in allelopathic interactions (Anaya et al., 1990). RGs are also responsible for the purgative properties of powders of *Ipomoea purga* root – which accumulate resin glycosides up to 20% of their root dry weight – and which are used in traditional Indian and Mexican medicine (Eich, 2008; Pereda-Miranda et al., 2010).

Despite their structural and functional characterization in multiple species, the diversity of RGs has not yet been assessed in a phylogenetic context. In addition, due to incomplete sampling, it is not clear if the characterized RG structures from different species are representative only of specific species or are broadly conserved across the entire family. Most structures have also been characterized using NMR spectroscopy, which is not a high-throughput method of assessing overall phytochemical diversity. Characterizing common patterns of fragmentation in mass spectrometry can help in the rapid identification of RGs.

To address these goals, we sampled 26 Convolvulaceae species using ultra high-performance liquid chromatography tandem mass spectrometry (UHPLC-MS/MS), characterized the structural diversity RGs in a phylogenetic context, and structurally elucidated a novel RG in the species *Dichondra argentea*. This study was enabled by our preliminary observations that RGs tend to fragment in a characteristic manner producing some signature fragments. We built upon this observation by computationally sampling a wide repertoire of possible moieties in the RG structural space. Our investigation revealed thousands of LC-MS peaks with MS/MS fragmentation patterns similar to known RGs but with unknown structures. This is emblematic of most LC-MS/MS studies of natural extracts where a majority of the peaks cannot be annotated. Thus, in this study, we investigated the RG fragmentation patterns in greater detail and developed a computational pipeline for identifying the putative structural components of over 1600 RG peaks, most of which are novel. This study lays the foundation for further molecular analysis of Convolvulaceae RGs.

## Results

### Sampling and identification of resin glycosides from Convolvulaceae species using MS/MS and NMR

To characterize RG structural diversity, we first generated a collection of 31 plants from 26 Convolvulaceae species (**Supplementary File 1**). Six of the twelve tribes and all five major clades in the family (Stefanović et al., 2003, 2002) were represented. The identity of each of these species was verified by sequencing and comparative analysis of a part of the *maturase K* (*matK*) gene region from the plastid DNA **(Supplementary File 2)**. We found that while most species were correctly defined, some such as *Distimake quinquefolius*, *Ipomoea pubescens* and *Calystegia sepium* needed re-assignment as *Ipomoea sp.*, *Merremia sp.*, and *Convolvulus sp.*, respectively **(Supplementary Fig. 1).**

Root and leaf extracts of these 31 plants, grown in growth chambers and greenhouses, were obtained at variable stages of growth and assessed using UHPLC-MS/MS **(Supplementary File 3)**. The goal of this analysis was to obtain a snapshot of the RG profiles of individual species, similar to the profiling performed for Solanaceae-type acylsugars from the New York Botanical Gardens (Moghe et al., 2017). Thus, consistency of growth stage and conditions – which is impossible when sampling 26 globally distributed species with variable growth habits and evolutionary adaptations to different environments – was not maintained. Exploratory analysis revealed substantial profile diversity for jalapinolic acid (11-hydroxyhexadecanoic acid; C16-OH; m/z=271.2279) containing compounds ( **Fig. 2A,B; Supplementary Fig. 2A–D)**, which may indicate species-specific RG patterns.

**Figure 2:**
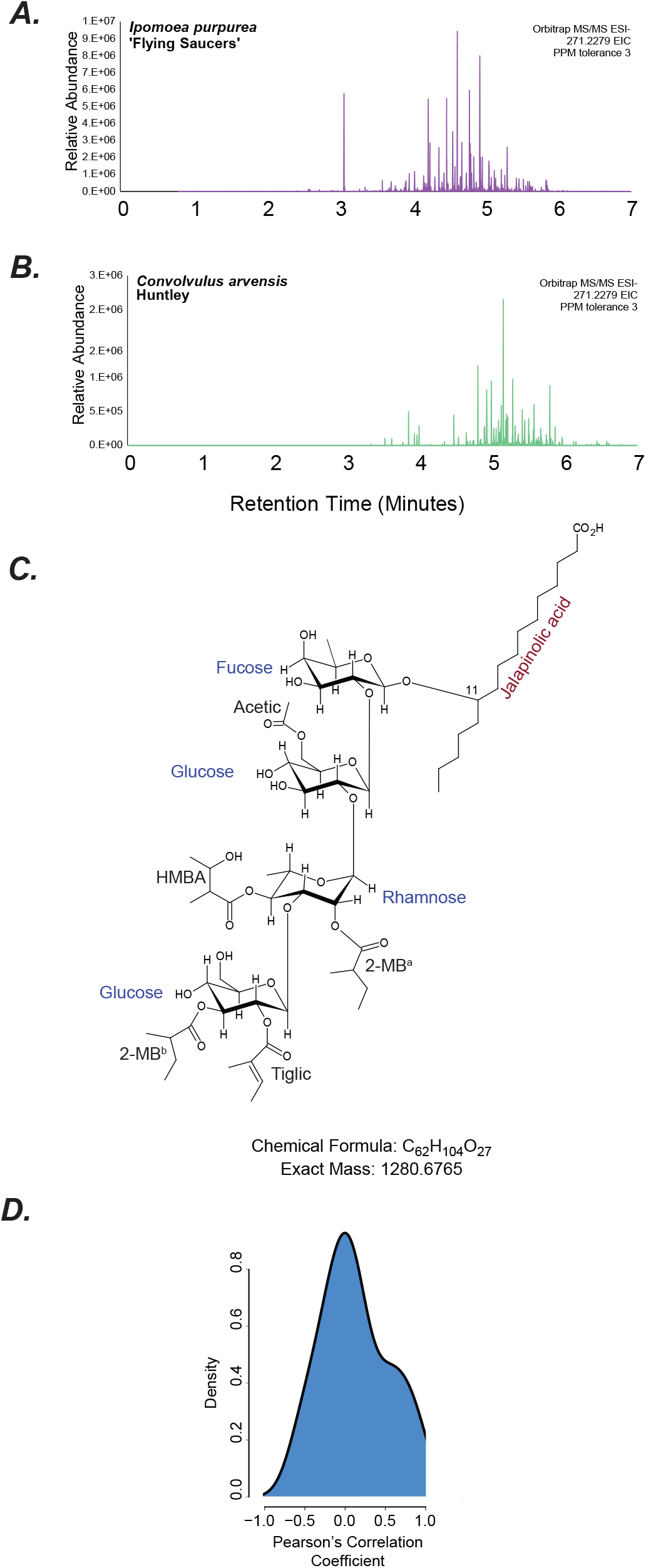
Chromatograms of resin glycoside peaks from different species. (A,B) Extracted ion chromatograms of m/z 271.2279 -- corresponding to the mass of jalapinolic acid -- from *Ipomoea purpurea* and *Convolvulus arvensis*. (C) NMR-inferred structure of Dichondrin D extracted from *D. argentea* Silver Falls. (D) Distribution of PCCs of relative root and leaf RG peak areas calculated in each species.

After pre-processing and filtering the peaks obtained using UHPLC-MS/MS, we computationally identified high-confidence RG-like peaks from the untargeted metabolomics data using multiple criteria **(Supplementary Fig. 2E)**. The primary criterion for RG peak selection was the presence of fragment pairs – one, a hydroxy- or dihydroxy-acyl chain of varying lengths, and two, the acyl chain attached to a sugar (pentose, deoxyhexose or hexose) **(Supplementary File 4, see Methods)** – defined based on previous knowledge (Eich, 2008; Pereda-Miranda et al., 2010) and our empirical observations **(Supplementary Fig. 3,4).** Overall, 2687 peaks congruent with the above two criteria and other filtering criteria were identified across all sampled species **(Supplementary File 1)**. We randomly and manually assessed multiple observable peaks containing the fragment pairs in different samples and found that most or all of the peaks were being captured by our computational pipeline. However, we cannot completely rule out the possibility that using these signature fragments underestimated the RG diversity in our samples.

To confirm whether the peaks identified were indeed RGs, we catalogued 30 previously characterized RGs from 6 species in our study **(Supplementary File 5)** and asked whether they were detected in our data. Thirteen out of the 30 RGs were captured in our processed dataset, and the rest were not detected. As seen also in later results, the low number of detected RGs likely speaks to the within-species RG structural diversity. None of the eight previously identified *C. arvensis* RGs we detected in our study, despite 150 and 215 RG signals detected in the two *C. arvensis* accessions sampled **(Fig. 3)**.

**Figure 3:**
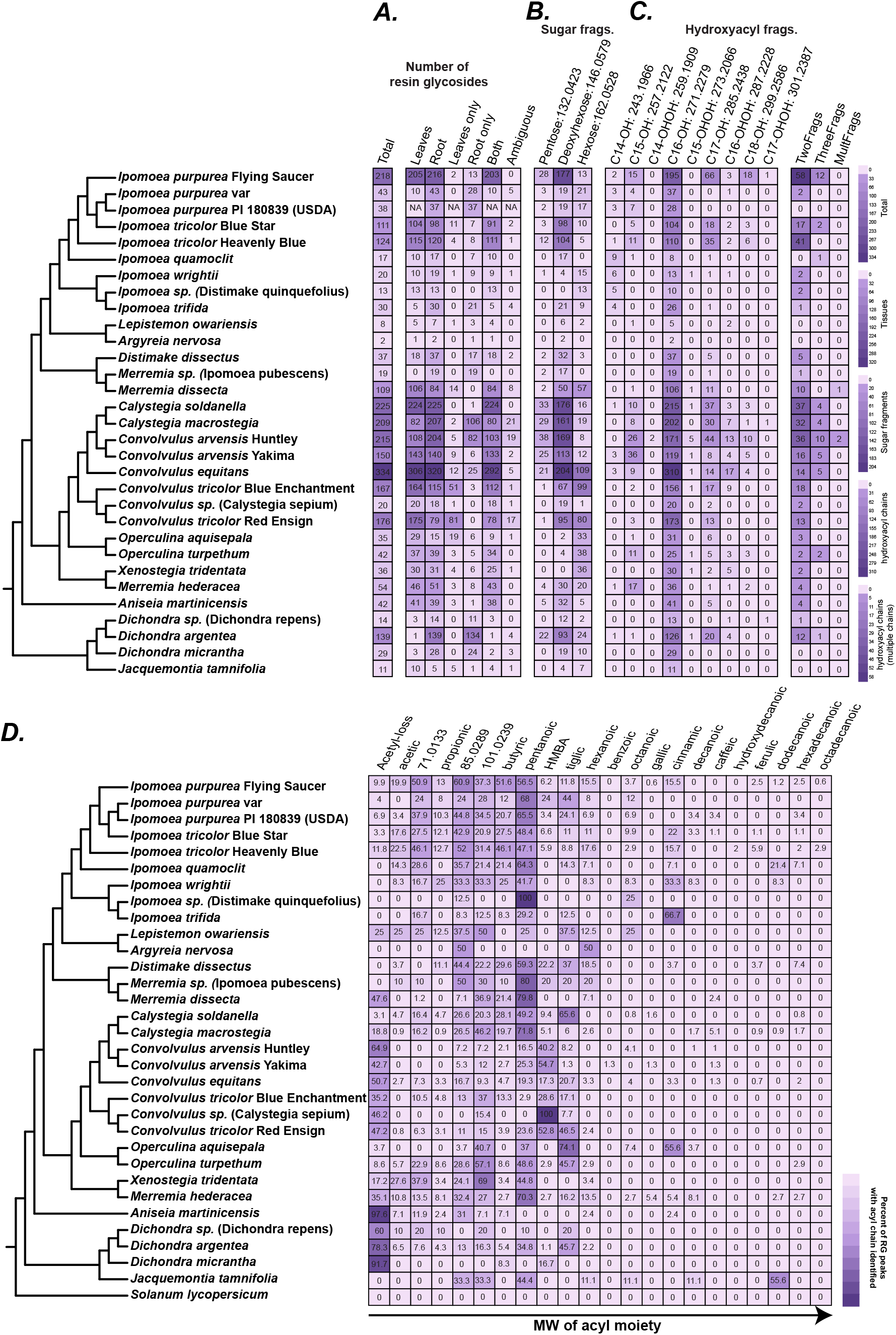
Phylogenetic, organ-wise and structural diversity of resin glycosides. In the phylogenetic tree, the name in brackets is the original name on the seed stock, and the name in italics is the predicted species/genus based on matK phylogeny. (A) Number of RG peaks split by their organ of occurrence. See Methods for details on thresholds used for these definitions. (B) Number of RG peaks with neutral loss corresponding to pentose, deoxyhexose, or hexose groups typically seen attached to the hydroxyacyl chain involved with macrolactone ring formation (C) Number of RG peaks with MS/MS fragments corresponding to the estimated masses of the different hydroxy- and dihydroxyacyl fragments. Some RG peaks had more than one fragments, as shown in the last three columns. (D) Percentage of RG peaks having a given acyl chain out of all RG peaks in that species with an acyl chain prediction. Numbers do not add up to 100 since each RG may have multiple acyl chain predictions. The percentages provide an idea of the general trend of acyl chain usage in a given species. Inference of the acetic acid group is based on neutral loss while other inferences are based on observation of specific MS/MS fragments. Monoisotopic [M-H]-acyl masses that cannot be confidently assigned to known acyl chain fragments are noted.

To further confirm the detection of RGs, we purified and structurally elucidated one highly abundant predicted RG peak with [M-H]-m/z 1279.67, from *Dichondra argentea* Silver Falls using NMR. Its molecular formula was determined to be C_62_H_104_O_27_, with the structure comprising a fucose, two glucoses and a rhamnose **(Fig. 2C, Supplementary Table 1; Supplementary Fig. 5)**. The jalapinolic acid moiety was found to be in an open structure instead of in a macrolactone ring. Previous RGs identified from *Dichondra repens* were also found to contain two rhamnoses and glucoses (Dichondrin A, B) or two rhamnoses and one glucose (Dichondrin C) (Song et al., 2015), both with open hydroxyacyl chain instead of a macrolactone ring. We thus call this structure Dichondrin D. The MS/MS fragmentation pattern of Dichondrin D could be explained by the characterized structure **(Supplementary Fig. 6)**. This result further confirmed that the fragmentation patterns being assessed were from true RGs.

### Resin glycoside accumulation patterns are generally consistent within a single plant but vary between varieties and species

Substantial previously uncharacterized RG diversity was found at various phylogenetic scales **(Fig. 3)**. First, we found RGs in all sampled species including smaller clades such as *Dichondra* and *Jacquemontia* (Stefanović et al., 2002). RGs have also been documented in *Cuscuta* (Du et al., 1998), whose phylogenetic placement in Convolvulaceae is not completely clear (Stefanović and Olmstead, 2004). This suggests that the evolution of RGs occurred in the ancestor of all Convolvulaceae species. Second, at the genus level, over a hundred and up to 334 RG signals were detected from different *Convolvulus*, *Calystegia*, *Ipomoea*, *Dichondra* and *Merremia* species. In contrast, other genera showed only 1-40 RGs. While this could indicate low RG diversity in those species under the sampling conditions, sampling other species in those genera could reveal additional diversity. Substantial variability was also found at the species and variety levels. Over 300 RG signals were found in *Convolvulus equitans* – the most among all plants sampled, while an unidentified *Convolvulus*/*Calystegia* species produced only 20 RGs. Two *Convolvulus arvensis* accessions collected from Yakima and Huntley in Washington state, USA that are ~200 miles apart produced 150 and 215 RGs, indicating genotype, environment or genotype x environment interactions in defining RG profiles. The fact that only 13/30 previously identified RGs could be detected in our study from the same species **(Supplementary File 5)** further supports this inference.

Metabolite sampling from the 26 species was performed from both leaves and roots, providing an opportunity to understand organ-specific localization of RGs. We divided the signals into three groups – only leaves, only roots, and both **(Fig. 3A)**. In almost all cases, RG signals were found in both organs. Analysis of RG peak areas in individual species from leaves and roots revealed low correlation between the peak areas in almost all species **(Fig. 2F; Supplementary File 1)**. The most abundant RGs in roots vs. leaves were also very different between roots and leaves of every species **(Supplementary File 1)**. In *Merremia sp.*, three *Dichondra* species and two *Ipomoea* species (*I. trifida*, *I. purpurea*), most or all RGs were found in the roots. This observation strongly suggests that RGs are produced in the roots. It is not clear, however, whether RGs are also produced in the leaves or are simply transported there.

### Substantial structural variation exists among resin glycosides

The phylogeny-guided sampling of RGs revealed substantial structural diversity in each species at the level of oligosaccharide lengths, hydroxyacyl chains, and the decorating acyl chains. Previous studies have reported existence of hydroxyacyl chains of varying lengths; thus, we computationally scanned the identified RG peaks to identify the type of long acyl chains seen in our data. Overwhelmingly, we saw m/z=271.2279, which corresponds to 11-hydroxyhexadecanoic acid (jalapinolic acid) **(Fig. 3B)**. The most diversity in hydroxyacyl chains was found, unsurprisingly, in the two species with the most overall RG diversity – *C. arvensis* and *C. equitans* – with hydroxypentadecanoic acid (C15-OH; m/z=257.2122), dihydroxyhexadecanoic acid (C16-OHOH; m/z=287.2228) and hydroxyoctadecanoic acid (C18-OH; m/z=299.2586) found in multiple RGs.

The MS/MS peak m/z=271.2279 co-occurs with m/z=417.2854 **(Supplementary Fig. 4)**, which corresponds to C16-OH bonded to a deoxyhexose. We thus searched the MS/MS data across all species to identify hydroxyacyl chains of all lengths with the three types of sugars previously reported in RGs, namely pentoses (xylose), deoxyhexoses (rhamnose, fucose, quinovose) and hexoses (glucose) in RGs **(Fig. 3B)**. Most RG peaks had the above noted m/z peak association, however, a substantial number were also associated with a hexose, which most likely is glucose. The relative frequencies of pentose vs. deoxyhexose vs. hexose was species-dependent. Many *Convolvulus* and *Calystegia* species produced >20 RGs with deoxyhexose and pentose as the most and second-most frequent sugars, respectively, while the *Merremia* and *Ipomoea* species contained deoxyhexose and hexose **(Fig. 3B)**. Previously identified RGs from *Calystegia* show the hydroxyacyl macrolactone ring connected to a glucose (a hexose) and quinovose (a deoxyhexose) (Gaspar, 1999; Takigawa et al., 2011). Since the sugar-hydroxyacyl paired fragment in our study detects only one end of the macrolactone ring connection, the glycosidic linkage between deoxyhexose and the hydroxyacyl chain may be more stable than the ester linkage with hexose in electrospray ionization and collision-induced dissociation in LC-MS. Both pentose and hexose were equally frequent in *D. argentea* as an alternative sugar **(Fig. 3B)**. In contrast, the two *Operculina* species and *Xenostegia tridentata* had hexose instead of deoxyhexose or pentose as their most preferred sugar for adding the hydroxyacyl chain. Eight previously published structures from *O. macrocarpa* and *O. turpethum* (Ding et al., 2012; Lira-Ricárdez et al., 2019) with the hydroxyacyl chain bonded to glucose agree with this finding. Based on these observations, we infer that hydroxyacyl chain addition (or at least the glycosidic bond formation) with deoxyhexoses was likely the ancestral state for RG biosynthesis, with lineage-specific shifts to hexose or pentose. The mechanism of this addition is not clear.

We also tested the occurrence of fragments corresponding to small aliphatic or aromatic acyl chains esterified to the sugar hydroxyl groups **(Fig. 3D)**. Using a database of all possible acyl CoAs from the Chemical Entities of Biological Interest (ChEBI) database (Degtyarenko et al., 2008) between 50-200 Da (the mass range of Solanaceae-type acylsugar acyl groups) as well as previously identified RG acyl groups (Eich, 2008; Pereda-Miranda et al., 2010), we asked if the fragment masses of the small acyl chains could be identified. Twenty-two acyl-related fragment ion exact masses were searched for **(Supplementary File 4).** The most frequent m/z seen across multiple RG peaks was m/z=101.061, corresponding to five carbon aliphatic saturated acids (e.g. methylbutyric, isovaleric, pentanoic) **(Fig. 3D)**, which we cumulatively refer to as C5 or “pentanoic” below. Mass-to-charge ratios of 71.0133, 85.0289 and 101.0239 normally correspond to acrylic, crotonic and acetoacetic acids respectively, however, it is not clear if the occurrence of these fragments indicates presence of these chains rarely found in previous RGs or whether they are degradation products of larger groups. For example, sequential demethylation and unsaturation of a C5 chain from in-source fragmentation can also produce fragments of m/z 85.0289 and 71.0133.

Differences in acyl incorporation between species were observed. RGs from *Convolvulus* species tended to contain m/z=117.059 (hydroxymethylbutyric acid, HMBA) while those from *Ipomoea* had m/z=101.061 (pentanoic acid) **(Fig. 3D; Supplementary Fig. 7)**. Cinnamic acid was predicted mostly in *Ipomoea* species; presence of *trans*-cinnamic acid has been previously reported in Ipomoeas (Noda et al., 1992; Barnes et al., 2003; Castañeda-Gómez and Pereda-Miranda, 2011). *Calystegia* species were all predicted to contain C5 acyls, except *C. soldanella* also with substantial tiglic RGs. This pattern is corroborated by studies in *C. hederacea* and *C. soldanella* containing 2*S*-methylbutyric acid as the C5 acyl chain (Ono et al., 2021; Takigawa et al., 2011). As per prediction here, the two studies in *Operculina* also demonstrate a dominance of C5 (2*S*-methylbutyric) and tiglic acids in two *Operculina* species (Ding et al., 2012; Lira-Ricárdez et al., 2019). Generally, short chain fatty acids were more represented in RG structures than long chain saturated and unsaturated fatty acids.

### Molecular networking reveals significant structural differentiation between *Convolvulus* and *Ipomoea* resin glycosides

To visualize the structural differences in RGs across the Convolvulaceae, we clustered MS/MS fragmentation patterns using molecular networking, which groups similar spectra based on their cosine scores. A total of 2100 RGs (78.1% of detected RGs) were clustered into a single large network, and these were displayed using genus-level occurrence patterns **(Fig. 4)**. We found large, centrally located regions 1 and 2, comprising fragmentation spectra shared by multiple genera. The greatest difference was seen between *Ipomoea* and *Convolvulus* – although these genera shared peaks in regions 1 and 2, dozens of unique RGs were found in regions 4,6,7 (*Convolvulus*) and region 3 (*Ipomoea*). These observations suggest fixed lineage-specific differences between the major genera. Analysis of the pseudo-molecular ion masses **(Fig. 4B)** revealed that *Convolvulus* produced hundreds of high molecular RGs with m/z>1500 while *Ipomoea* species produced many low molecular weight RGs not observed in other species. This finding agrees with previously reported RGs from *Convolvulus arvensis*, containing many penta-, hexa- and hepta-saccharides (Fan et al., 2018; Lu et al., 2021) vs. predominantly tri- and tetra-saccharides from *Ipomoea* species (Cruz-Morales et al., 2016; Pereda-Miranda et al., 1993). Above results **(Fig. 3D)** showing differential acyl chain patterns in *Ipomoea* vs. *Convolvulus* may also contribute to overall spectral divergence.

**Figure 4:**
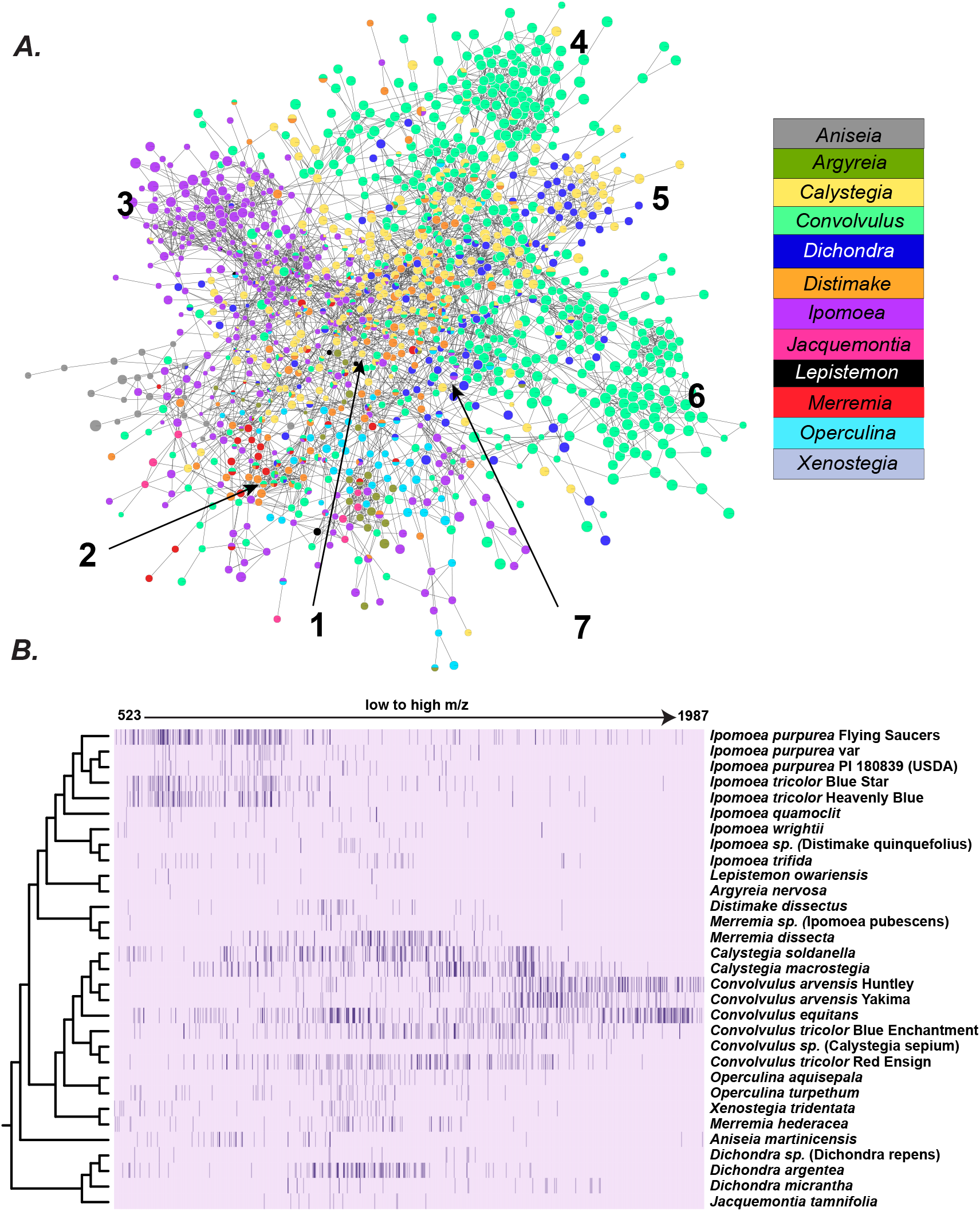
Structural diversity of resin glycosides across all sampled species. (A) MS/MS molecular networking shows shared and divergent structural diversity of RGs. The graph is loosely classified into 7 regions based on manual observation of the distinct regions. (B) In the heatmap, columns are representative of MS1 masses of RG peaks found across all sampled species organized from low to high m/z. The rows are the species. Each cell is colored by how many peaks of a given m/z are detected in a given species. Typically, the values in the cells correspond to 0,1 and 2.

### A knowledge-based algorithm can explain many MS/MS peaks and help predict resin glycoside structural components

Based on the above substructures identified across our entire LC-MS/MS dataset, we sought to rapidly annotate the MS/MS fragments of individual RG peaks. The Metabolomics Standards Initiatives guidelines (Sumner et al., 2007) specify four levels of annotation, with Level 3 corresponding to putatively identified compound classes, and Level 2 corresponding to putatively identified compounds. The compounds called RGs herein are annotated at Level 3 based on MS/MS information and validating NMR data from one peak. Acquiring Level 2 annotation for individual peaks requires information about the types and stereochemistry of sugars and acyl chains that is not available from LC-MS/MS experiments. However, the spectral information can be used to predict the structural components of the identified RG peaks, and thus narrow their structural neighborhood.

To facilitate putative component identification, we first attempted to better understand the different types of peaks produced upon fragmentation and any patterns of their co-occurrence. Three types of peaks showed frequent co-occurrences: (i) the hydroxyacyl fragment + one and two sugar (typically, 271–417–561/579), (ii) formate adduct (−46) or CO_2_ (−44) losses paired with a specific MS/MS fragment, and (iii) water (−18) loss **(Fig. 5)**. This water loss could arise from an open or closed macrolactone ring structure – both forms are found in diverse RG structures across Convolvulaceae and in our NMR results **(Fig. 2)** – or via a dehydration-led rearrangement occurring during electrospray ionization (Demarque et al., 2016) **(Fig. 5)**.

**Figure 5:**
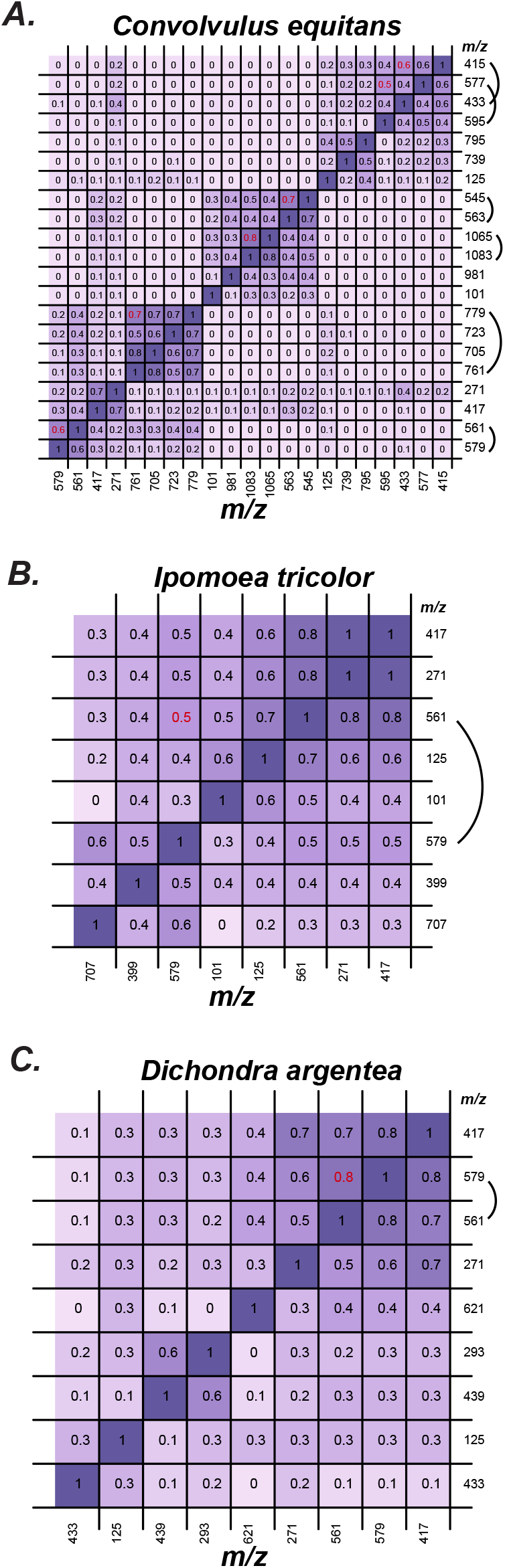
Correlations between MS/MS fragments. Pairwise Pearson’s Correlation Coefficient between nominal masses shown here. Masses highlighted by a curved arc indicate correlated pairs with a loss of 18, indicating the two fragments are closely associated with each other (correlation value in red). Possible explanations for the loss of 18 are shown in Supplementary Fig. 8.

With the above spectral knowledge, we established a pipeline for RG component prediction. First, we matched the top 20 observed MS/MS peaks of each MS1 peak to an in-house database of known moieties found in RGs (Eich, 2008; Pereda-Miranda et al., 2010) **(Supplementary File 4)**. Second, we performed pairwise comparisons of all fragments, and inferred neutral losses and the potential moieties those losses would correspond to. On an average, 72.5% of the 43,261 total MS/MS fragments across all samples could be annotated automatically using this manner. Third, the difference between the maximum MS/MS fragment with annotation and the pseudo-molecular ion mass of the parent compound was used to infer the remaining possible moieties. To reduce false positive predictions, we only considered acyl chains with occurrence in >5% of the RG peaks (“Frequent”**, Supplementary File 4)** and only predicted up to hepta-saccharide cores with up to three frequent acyl chains. We also considered water loss and formate/no-formate adducts. Using these rules, 1245 (45.9%) of the RGs were assigned putative component identifications **(Supplementary File 6)**. Multiple predictions were also obtained for each structure (low precision), all of which we provide for further manual consideration. Using knowledge of observed MS/MS fragments, prediction of formate-related and other neutral losses can help filter through the multiple predictions manually and thereby obtain a smaller set of component predictions.

Of the 13 RGs identified in our analysis, correct prediction was obtained for 11 of them (high recall; **Supplementary File 5**), with Dichondrin D **(Fig. 2C)** containing five acyl chains and Merremin D containing decanoic acid being the exception. When the algorithm was extended to five acyl chains and included decanoic acid, correct prediction was obtained for both RGs. As an example of the prediction, Tricolorin A is a tetra-saccharide containing two rhamnoses, one fucose, one glucose, 11-hydroxyhexadecanoic acid and two methylbutyric acid groups. There were two additional peaks corresponding to the m/z of its formate adduct 1067.563, which could correspond to Tricolorins E and D – structural isomers of Tricolorin A **(Fig. 6A)**. These peaks were accurately predicted to contain three deoxyhexoses and one hexose, two C5 acyls and a C16-OH. MS/MS peaks for multiple moieties were correctly isolated **(Fig. 6B)**.

**Figure 6:**
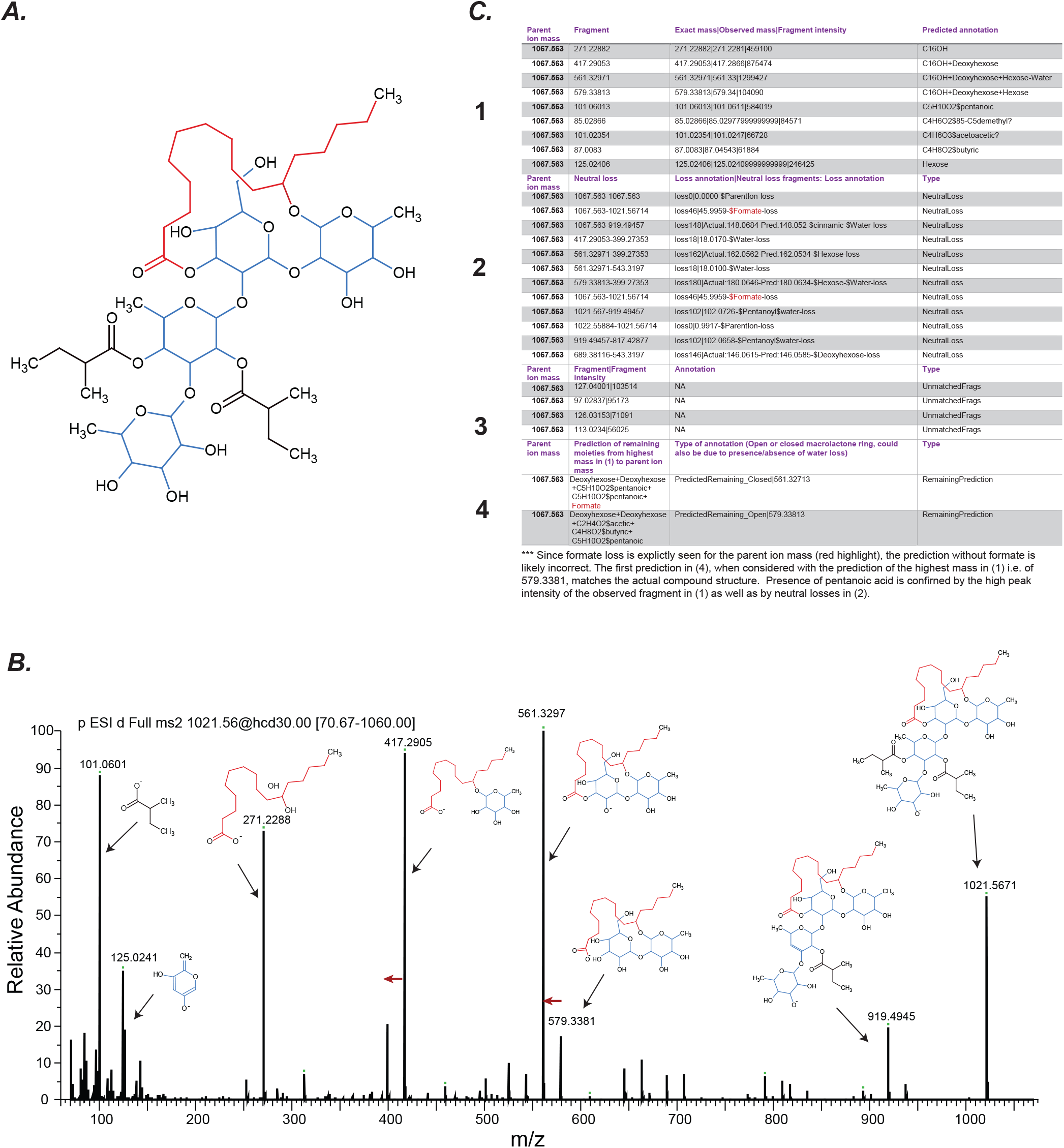
Example of component prediction using a knowledge-based approach. (A) Non-isomeric structure of tricolorins (e.g. tricolorin A). (B) Observed MS/MS fragmentation pattern of the most abundant tricolorin in *Ipomoea tricolor* Heavenly Blue. Red arrows show water loss. (C) Components predicted by comparing the observed MS/MS fragments to database masses. Maximum top 20 fragments based on peak intensity are chosen. Neutral losses are predicted by computing pairwise mass differences between the fragments and comparing them to database masses. The final structure is the combination of the components assigned to the highest mass in (1) and the most likely component combination in (4). Studying (2) and (3) fragments can help confirm or select appropriate predictions from (4).

Although the recall of this knowledge-based approach is high, the precision is low. One reason is presence of some ions with insufficient good quality fragmentation to produce informative peaks. The most common issue however is multiple predictions due to multiple residue combinations producing the same exact mass **(Supplementary File 6)**. For example, an RG containing a deoxyhexose and pentanoic acid will have its mass equivalent to one with hexose and hydroxymethylbutyric acid – both being common side chains. This issue can be mitigated by manually checking the extracted MS/MS fragments for presence/absence of formate adduct and specific acyl chains **(Fig. 6C)**. Nonetheless, the computational pipeline is able to explain a large proportion of the top MS/MS fragments of RGs and is able to provide tentative identifications of the RG structure based on the universe of likely fragments. Future iterations of the workflow can potentially allow for more precise predictions based on species-level component preferences.

## Discussion

In this study, we performed a phylogeny-guided analysis of RG structural diversity. Substantial diversity was found in RG structures in each plant, within species as well as between species. All species were found to accumulate RGs in the roots. Evidence of five different hydroxyacyl groups complexed with sugars with occurrence in >10 RGs was obtained, with C16-OH being the most common **(Fig. 3C)**. This was followed by hydroxyacyl chains of C15, C17 length, and then by C16-OHOH. Odd-chain fatty acids (C15, C17) are rarer compared to even-chain ones (Park and Nicaud, 2020), however, both have previously been described in RGs (Ding et al., 2012; Fan et al., 2018). In addition, hydroxylated C17 fatty acid is the second most prevalent macrolactone ring fatty acid except in *C. arvensis* Yakima, which preferred C15 as the alternate chain. Biosynthesis of odd-chain fatty acids has been proposed to occur (i) from propionyl coA as the primer in fatty acid elongation pathway instead of acetyl coA, (ii) via one carbon elongation of fatty acids e.g. in acylsugars (Kroumova and Wagner, 2003), and (iii) α oxidation of the more abundant even-chain fatty acids (Jansen and Wanders, 2006), but which of these mechanisms is prevalent in morning glories is unknown. These observations – combined with the unusual 11^th^ position hydroxylation in the hydroxyacyl chain – reveal the unique biochemistry of RGs. 12-hydroxyhexadecanoic acid and 11*(S)*-hydroxyheptadecanoic acid have been previously documented in *O. turpethum* (Ding et al., 2012) and *C. arvensis* (Fan et al., 2018), respectively but our study reveals the previously unreported extent of hydroxyheptadecanoic acid’s prevalence across the Convolvulaceae RGs. Furthermore, based on the chain distribution patterns **(Fig. 3C)**, we predict that a single enzyme - preferring jalapinolic acid and with promiscuous activities with other hydroxy/dihydroxy acyl chains – is responsible for acylation of the oligosaccharide core.

Among the acyl chains, five-carbon long chains (e.g., valeric acid, 2-methylbutyric acid, 3-methylbutyric acid) were the most frequent while four-carbon moieties (e.g., isobutyric acid, nbutyric acid), tiglic and acetic were also found in >10% of the RGs. Longer saturated acyl chains were not as frequently detected, indicating the predominant role of branched chain amino acid biosynthesis – from which most of the small acyl chains are likely derived – in RG decorations. Such diverse acylations are indicative of the involvement of BAHDs, similar to the promiscuous BAHDs of the Solanaceae acylsugar biosynthetic pathway (Fan et al., 2019; Kruse et al., 2020; Moghe et al., 2017). Solanaceae-type acylsugar BAHDs show conservation of acylation positions in orthologs despite their acceptor promiscuity, however, in RGs, analysis of previously characterized structures does not suggest positional uniformity (Eich, 2008; Pereda-Miranda et al., 2010). Thus, while we cannot completely rule out allelic divergence in a single fast-evolving BAHD as leading to the acylation diversification, it is more likely that multiple BAHDs play a role in RG decorations, and the loss/gain of BAHD activities in different lineages leads to the observed lineage-specific patterns of acylations.

Our findings reveal structural divergence indicative of underlying biosynthetic divergence in different lineages. At the genus level, the divergence between *Ipomoea* and *Convolvulus* is the most stark. Detailed analysis of spectral patterns performed in this study revealed some of the basis for this divergence – larger RGs in *Convolvulus* and use of different decorating acyl chains between *Convolvulus* and *Ipomoea* i.e., HMBA and pentanoic, respectively **(Fig. 3B)**. Identification of the underlying glycosyl and acyltransferases will help determine the molecular basis of this divergence.

The plant kingdom is estimated to contain over a million metabolites (Afendi et al., 2012), however, <5% can be confidently identified structurally using public databases (unpublished results). This inadequacy results in a vast proportion of the plant metabolome remaining unannotated, which can further create roadblocks in annotating the genes involved in metabolism. Here, we used the knowledge of fragmentation patterns in RGs to annotate their structural components. Such a knowledge-based approach is popularly used in lipid annotation by tools such as LipidBlast (Kind et al., 2013), whose internal database contains *in silico* fragmentation for >120,000 lipids from 30 structural classes. Recently, a deep learning method CANOPUS (Dührkop et al., 2020) was developed to predict metabolite classes using untargeted LC-MS/MS data. This approach was used to predict hundreds of flavonoids and anthocyanins in sweet potatoes (Bennett et al., 2021), however, more algorithmic development is needed for other phytochemical classes. Furthermore, much of the specialized metabolic diversity lies outside of current reference species that are the most well-studied (Moghe et al., 2017; Moghe and Kruse, 2018). Thus, approaches to rapidly annotate this “non-model diversity” are needed. The knowledge-based approach developed here for RGs provides a template for other acylsugars across the plant kingdom **(Fig. 1)**. Given the bonds remain generally the same – glycosidic, ester and ether linkages – we predict that the fragmentation pattern would be consistently reproducible if using comparable MS methodology. Overall, the findings presented here not only provide an extensive insight into the unique chemistry and diversification of RGs in Convolvulaceae – a horticulturally important family – but also provide a foundation for using the lineage-specific variation to probe the biochemistry of these compounds in the future.

## Materials and Methods

### Plant germination and growth conditions

Depending on the thickness of the seed coat, seeds were either sterilized using concentrated sulfuric acid or 10% trisodium phosphate (TSP) solution. Seeds with thick seed coats (e.g., *Ipomoea tricolor*) were acid scarified and sterilized with concentrated sulfuric acid for 15 minutes. Seeds with thinner seed coats (e.g., *Dichondra argentea*) were washed in 10% TSP for 10 minutes for surface sterilization. For both conditions, seeds were washed five times with sterile water afterwards. Student grade filter paper (VWR, Radnor, PA) was placed within a sterile petri dish and 1 mL of sterile water was added. The sterilized seeds were added to the petri dish and sealed with parafilm. Seeds were kept in the dark at room temperature for 4-10 days until germinated. When the radicle was clearly visible the seedlings were transferred to 4-inch pots and grown in a growth chamber under a constant light/dark (16 h/8 h) regime at 24 °C for 4 weeks. After 4 weeks, plants were moved to a greenhouse, re-potted if necessary and grown for approximately three months until samples were harvested.

### Genomic DNA extraction and amplification of maturase K (matK) partial gene sequences

Leaf samples were taken using clean tweezers and scalpel, and flash frozen in liquid nitrogen and stored at −80 °C until use. Approximately 50 mg of frozen tissue was ground in a 1.5 ml reaction tube using a disposable tube pestle. Genomic DNA from the ground leaves of all species grown was isolated using the E.Z.N.A. Plant DNA Kit (OMEGA Bio-Tek, Norcross, GA) following the manufacturers recommendations. The gDNA was eluted using 100 μl nuclease-free water and the concentration was estimated using a NanoDrop™ One instrument (ThermoFisher Scientific, Waltham, MA). The 1:10 diluted DNA was used to amplify a region of the maturase K gene using Q5® High-Fidelity DNA Polymerase (NEB, Ipswich, MA) following the manufacturers recommendations with following primer combinations: matK_F1 (5’-ATA CTT TAT TCG ATACAA ACT CCT TTT TTT-3’) and matK_R1 (5’-AGT ATT GCA ATT TGA ATA GTT TCA TTA C-3’) using 53 °C as annealing temperature or matK_F2 (5’-ATA CTT TAT TCG ATA CAA ACT CCT TTT TTT GGA AG-3’) and matK_R2 (5’-AGT ATT GCA ATT TGA ATA GTT TCA TTA CTC GAA A-3’) using 56 °C as annealing temperature. Both primer pairs amplify the same target while matK_F2 and matK_R2 bind approximately 40 bp upstream of their respective F1/R1 counterparts. PCR products were directly sequenced using Sanger sequencing at the Cornell Institute of Biotechnology using above primers.

### Confirmation of species identity via *matK* barcoding and generating a species phylogeny

The obtained *matK* sequences for each sampled species were searched against NCBI’s nr database using blastn (Camacho et al., 2009) default parameters to confirm that the correct species were sampled. All nucleotide sequences including those of the best hits were aligned using MAFFT v.7.453-with-extensions (Katoh et al., 2002) in Geneious Prime v.2020.1.1 using default parameters and provided as input to IQ-TREE v.1.6.10 (Nguyen et al., 2015), which was run with model selection (ModelFinder, (Kalyaanamoorthy et al., 2017)), single branch test using SH-like approximate likelihood ratio test (1000 replicates) (Guindon et al., 2010), and 1000 standard non-parametric bootstrap replicates using following parameters: -alrt 1000 -b 1000 -nt AUTO -ntmax 32. The optimal tree was obtained using the TVM+F+R2 model.

### Resin glycoside extraction

Leaves and roots of plants were harvested into 50 ml falcon tubes and snap frozen in liquid nitrogen. Frozen samples were rough ground and homogenized with a spatula. Approximately 100 mg of sample was weighed while keeping the material frozen with liquid nitrogen and 300 uL of methanol containing 10 uM telmisartan per 100 mg of tissue was added to samples (except in species 28-32; **Supplementary File 1**), and vortexed for 30 seconds. Two metal beads (2.3 mm) were added to the sample and samples were homogenized using a mixer mill (MM400, Retsch, Haan, Germany) at 30 bpm in 2-minute intervals for a total of 6 minutes within a 15-minute extraction period. Afterwards, samples were centrifuged for 30 seconds at 14,000 g to remove particulate. If supernatant still contained particulate, it was centrifuged again with the same parameters. Once all particulates were removed, the supernatant was transferred to HPLC vials for further analysis or stored at −80 °C.

### UHPLC-MS/MS run conditions

LC-MS analysis was performed on a ThermoScientific Dionex Ultimate 3000 HPLC equipped with an autosampler coupled to a ThermoScientific Q-Exactive Orbitrap mass spectrometer using solvent A (water + formic acid (0.1% v/v)) and solvent B (acetonitrile) at a flow rate of 0.6 ml/min, on a Phenomenex Kinetex 150mm C18 column (model: 00F-4462-AN). Metabolites were detected using full MS1 coupled to a data-dependent MS2 method, extracting the top 10 most abundant ions (Full-MS/ddMS2) in negative ionization mode. The total scan range used was 500 to 2000 m/z. Additional details of chromatographic, mass spectrometric and MS-DIAL-based data analysis methods are described in **Supplementary File 3**. RAW files from the LC-MS runs were converted to *abf* files using AbfConverter (Reifycs, Tokyo, Japan) and used for further data analysis with MS-DIAL (Tsugawa et al., 2015). Converted raw files for leaf and root samples for each species were aligned to each other, and an alignment-based mgf file was exported. These output files were then filtered using custom Python scripts as described in **Supplementary Fig. 2**.

### Liquid-chromatography mass-spectrometry (LC-MS) data analysis

RGs were identified using a multi-step analysis. First, we performed manual analysis of multiple LC-MS runs to study RG fragmentation patterns. In addition, previously reported MS/MS fragmentation patterns (Eich, 2008; Pereda-Miranda et al., 2010) as well as MS/MS fragmentation simulated in ChemDraw v19.0 (PerkinElmer, Waltham, MA, USA) provided us with a general understanding of the fragmentation pattern and allowed us to compile a database of RG-specific fragment ions **(Supplementary File 4)**. Automated identification and quantification of RGs in the analyzed samples was carried out using in-house Python scripts as outlined in **Supplementary Fig. 2E**.

### Identification of resin glycoside peaks from LC-MS data

The alignment files produced using MS-DIAL per species were analyzed individually. Multiple filters were applied using custom Python scripts to separate likely true RGs from potential noise, and the selected peaks were then extracted from the mgf format files. The filters applied were as follows: **(1)** The peak area of the selected MS1 peak in either leaf or root needed to be >10X the signal of that peak vs. Blank as well as >1000, **(2)** every selected peak had to contain at least one out of 30 special pairs of MS/MS fragments: Each pair comprised one hydroxyacyl fragment (out of 10) and a combined fragment of the hydroxyacyl bound to a sugar (out of 3) (e.g., 271.2279–417.2858 for deoxyhexose+C16-OH), **(3)** the MS/MS peak areas of the fragments needed to be >10% of the maximum MS/MS ion intensity, so as to reduce noise. The last filter also resulted in some true RG peaks being filtered out e.g., *m/z* 1079.4929 in *Operculina turpethum*, which contained MS/MS fragments 257.2108 and 419.2613 corresponding to a C15-OH complexed to a hexose sugar – a motif frequently found in this species. However, the 257.2108 fragment was only 4% of the maximum ion intensity. **(4)** MS-DIAL identifies peaks as linked to adducts or to ions of higher masses. While all adducts were processed as RGs, peaks linked to other peaks of higher masses were processed only to consider one of the two peaks. The Dichondrin D peak (1279.6695) was associated with a higher mass 1364.642, and thus was lost to downstream steps because of this filter. Distinction between only root, only leaves and ambiguous was made as follows: If the unnormalized peak area was >1000 in root or leaves and 0 in the other organ, it was assigned to root or leaves, respectively. On the other hand, if the peak area was >1000 in both, it was assigned to both organs. Peaks not satisfying these scenarios were deemed ambiguous.

### Compound purification for NMR

For structural elucidation of Dichondrin D, larger amounts were extracted from roots of *D. argentea.* Root tissues were collected and frozen in liquid nitrogen and stored at −80 °C until bulk extraction. First, the tissues were homogenized using a prechilled Victoria™ cast iron grain mill under constant infeed of liquid nitrogen to keep tissues frozen. Subsequently, 3 g of tissue were extracted using 9 mL of methanol for *D. argentea*. The extraction was performed under constant shaking at 25 °C for 15 minutes in an Eppendorf New Brunswick Excella E24 Orbital Shaker (180 rpm). Afterwards, the samples were vacuum filtered using a Büchner funnel and Whatman grade 1 qualitative filter paper to remove particulate matter. The extract was dried down until solidified using a rotary evaporator (Rotavapor R-II, Büchi, Flawil, Switzerland) equipped with a water bath set to 25 °C and a vacuum of 50 mbar. The solid samples were then reconstituted in 300 μL MeOH (HPLC grade). Dichondrin D was then purified on an Agilent HPLC 1100 System using an Agilent Eclipse XDB-C18 Semi-Prep (5μm, 9.4×250mm) column over 32 minutes at a flow rate of 3.5 mL per minute. The gradient started with 40% solvent A (acetonitrile, HPLC grade) and 60% solvent B (H2O with 0.1% formic acid, HPLC grade) from minute 0 to 3, and increasing the concentration of solvent A to 100% at minute 20, 100% solvent A for 5 minutes, followed by equilibrating the column to the starting conditions for 7 minutes. The samples were purified over 10 injections of 30 μL each in order to purify high enough amounts for subsequent NMR spectroscopy. Eluting RGs were examined using a Waters Quattro Ultima Triple Quad Micromass mass spectrometer and the corresponding fractions for Dichondrin D (*m/z* 1279.6, RT 19.5 – 20.0 min) were collected using a ISCO Foxy 200 Fraction Collector. The samples were further assessed using a ThermoScientific Dionex Ultimate 3000 HPLC coupled to a ThermoScientific Q-Exactive HF Orbitrap and determined to be of approximately 90% purity. The fractions of interest were then dried via rotary evaporation and reconstituted in 99.5% deuterated MeOH. The drying procedure was repeated, and the sample was reconstituted in 1 mL 99.9% deuterated MeOH before transferred to the NMR tube.

### NMR spectroscopy data acquisition and analysis

NMR spectra were recorded on the Bruker AVANCE III HD 800 MHz (800 MHz) spectrometer at State University of New York Environmental Science and Forestry NMR facility. 1H NMR chemical shifts are reported in ppm (δ) relative to residual solvent peaks (3.31 ppm for methanol-*d_4_*). ^13^C NMR chemical shifts are reported in ppm (δ) relative to residual solvent peaks (49.0 ppm for methanol). All NMR data processing was done using MNOVA 14.2.1 (https://mestrelab.com/).

### Molecular networking analysis

The mgf files and peak area files were submitted to GNPS (Wang et al., 2016) for molecular networking. The networking parameters were kept at default values except “Precursor Ion Mass Tolerance” was set to 0.05, “Min Pairs Cosine” was set to 0.5, “Network TopK” was set to 15, “Maximum Connected Component Size” was set to 0, “Minimum Matched Fragment Ions” was set to 3, and “Minimum Cluster Size” was set to 1. The resulting graphml file was imported into Cytoscape v 3.8.0 (Shannon et al., 2003) and visualized using the Prefuse Force Directed Layout.

### Data and code availability

Unfiltered mgf files as well as metadata is being deposited in MetaboLights under the accession ID MTBLS3113. All Python scripts for parsing MS-DIAL exported files and for generating component annotations have been deposited on the moghelab/ResinGlycoside repository. All uploaded files will be made publicly available upon manuscript acceptance.

## Acknowledgements

The authors would like to thank Dr. A. Daniel Jones at Michigan State University for helpful discussions on the topic, as well as Dr. Navid Mohaved for helping with LC-MS training and advice.

## Contributions

Performed experiments: AB, LK, EM, SJP, EL; Analyzed data: AB, LK, FS, GM; Wrote the manuscript: LK, GM, AB. All authors reviewed the manuscript.

## Supplementary Data

Supplementary Figures: 8

Supplementary Files: 6

Supplementary Table: 1

---

**Supplementary Figure 1:**
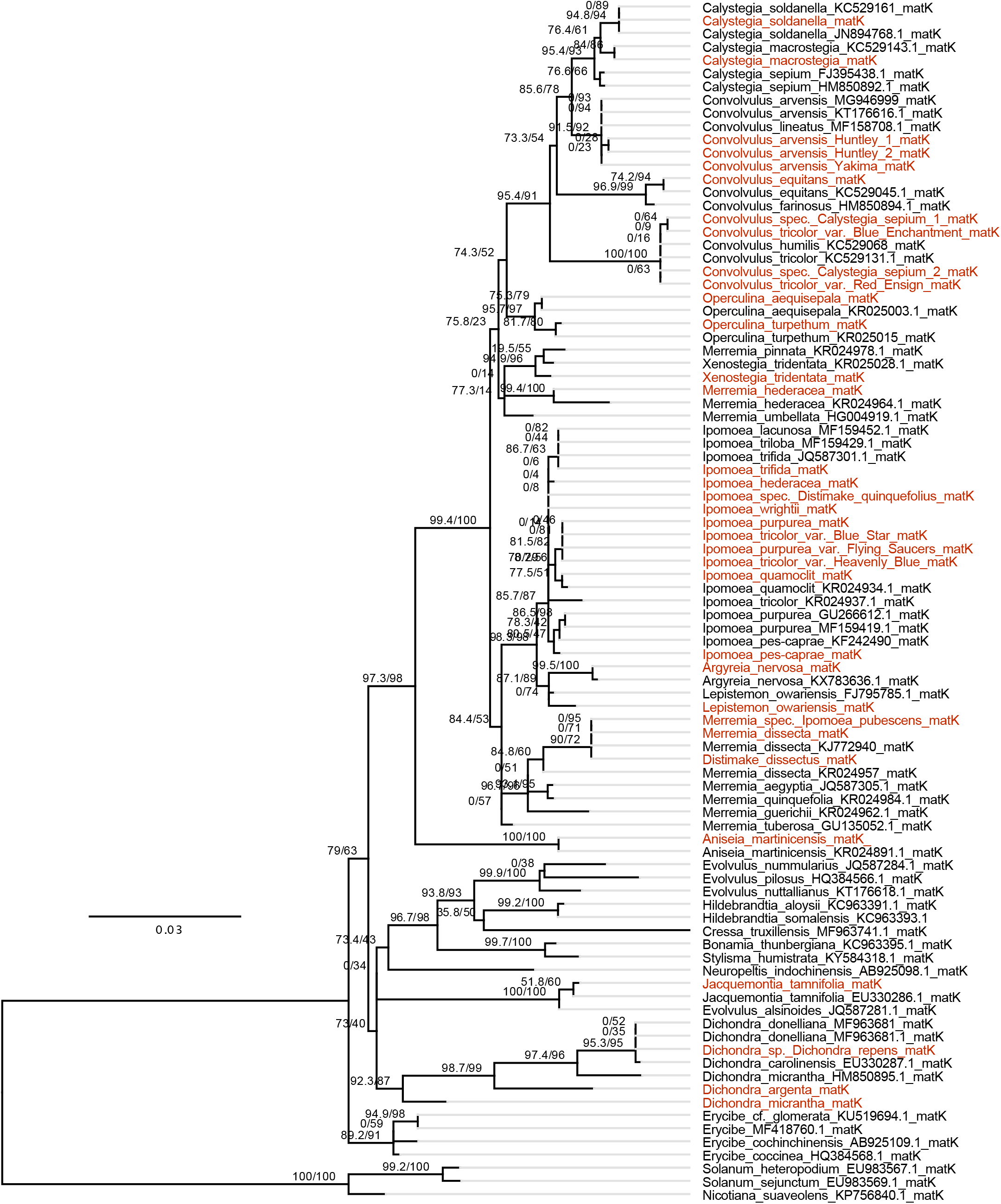
matK phylogeny of selected species. matK sequences were obtained from Sanger sequencing of the tested species (red) as well as from the NCBI database (black). The tree was optained with IQ-TREE using the SH-like approximate likelihood ratio test (SH-aLRT,1000 replicates) and 1000 standard non-parametric bootstrap replicates. Branch support values are SH-aLRT/bootstrap support. The tree is rooted on the brach leading to the included Solanaceae species.

**Supplementary Fig. 2:**
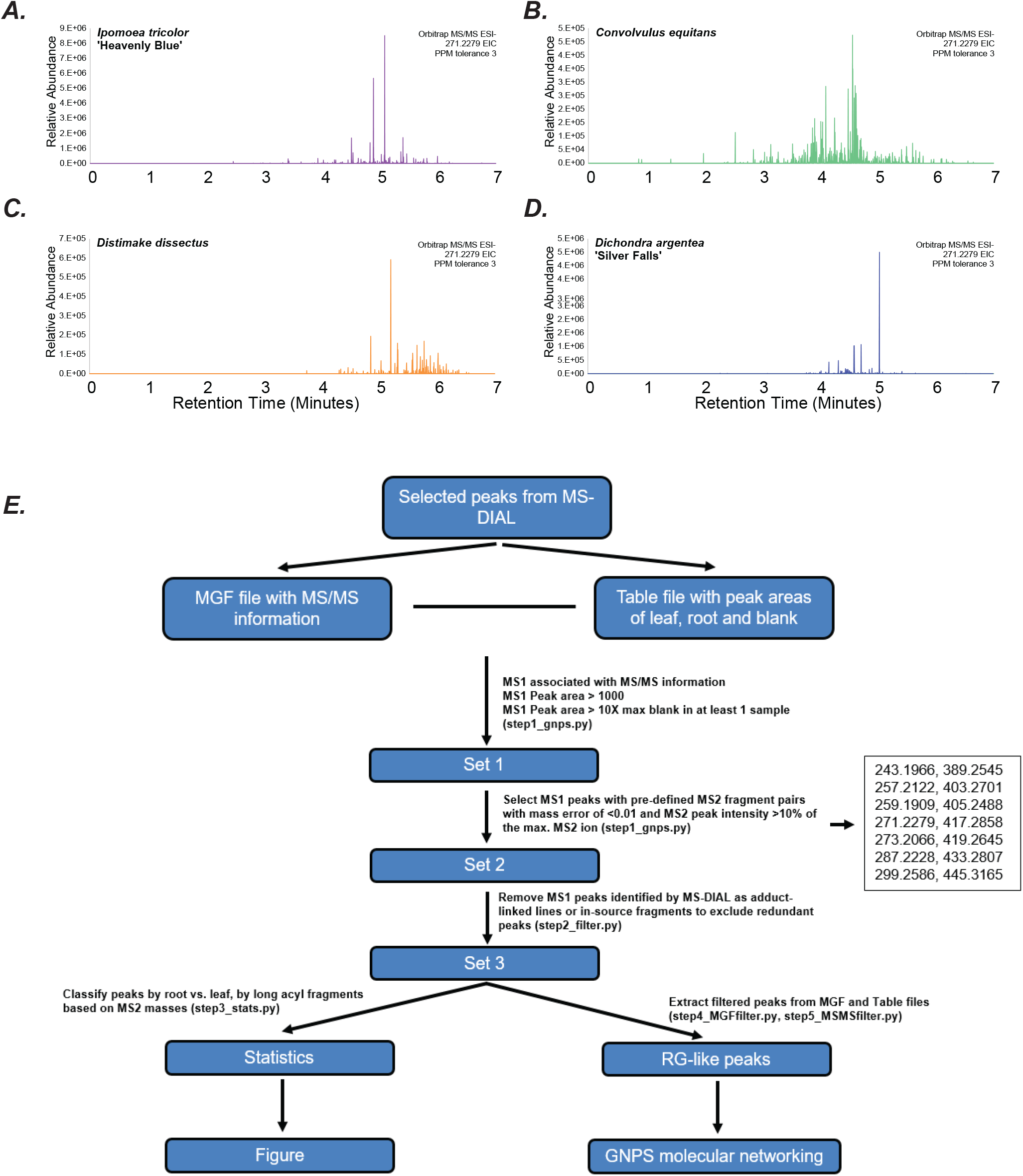
Resin glycoside diversity. (A-D) Extracted ion chromatograms of m/z 271.2279 -- corresponding to the mass of jalapinolic acid -- from four Convolvulaceae species. (E) Computational pipeline used for RG detection. An example of the RG signature pairs is shown, out of 30 pairs (10 acyl chains x 3 sugar types) used for actual analysis.

**Supplementary Figure 3:**
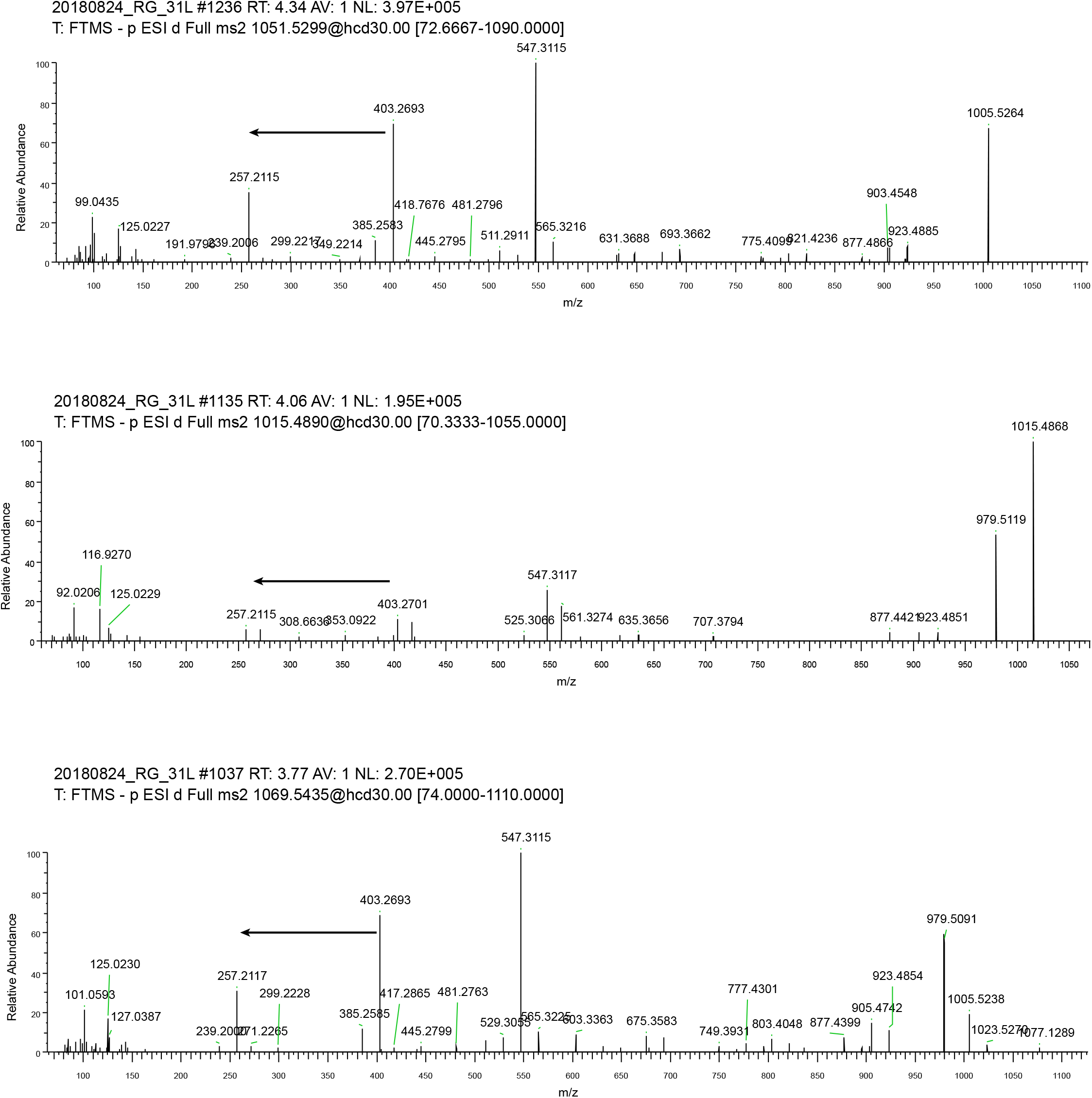
Paired MS/MS fragments for the 257 fragment. Arrows limit peaks corresponding to the hydroxyacyl chain (smaller mass peak) and the hydroxyacyl chain+sugar (larger mass peak).

**Supplementary Figure 4:**
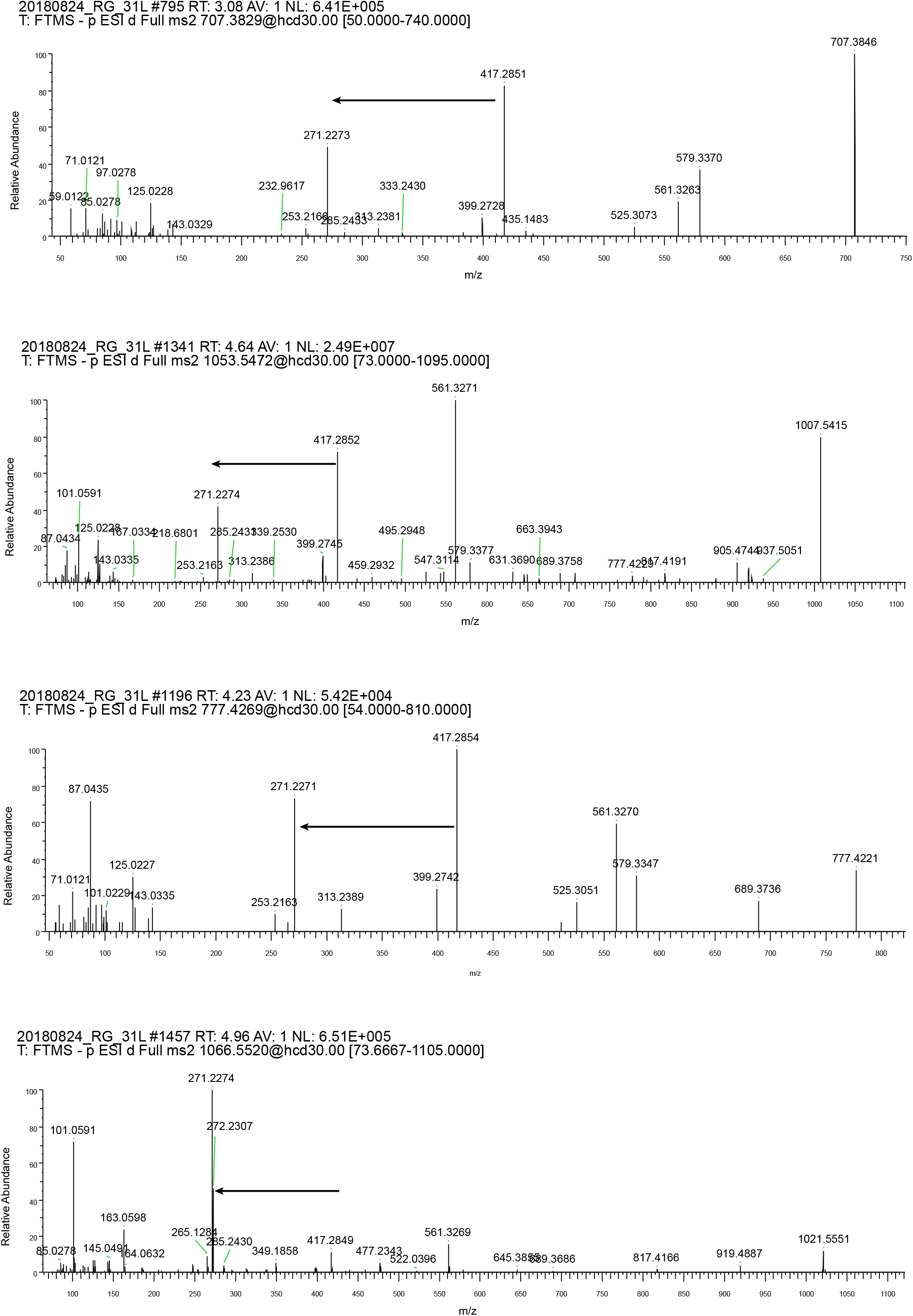
Paired MS/MS fragments for the 271 fragment. Arrows limit peaks corresponding to the hydroxyacyl chain (smaller mass peak) and the hydroxyacyl chain+sugar (larger mass peak).

**Supplementary Figure 5:**
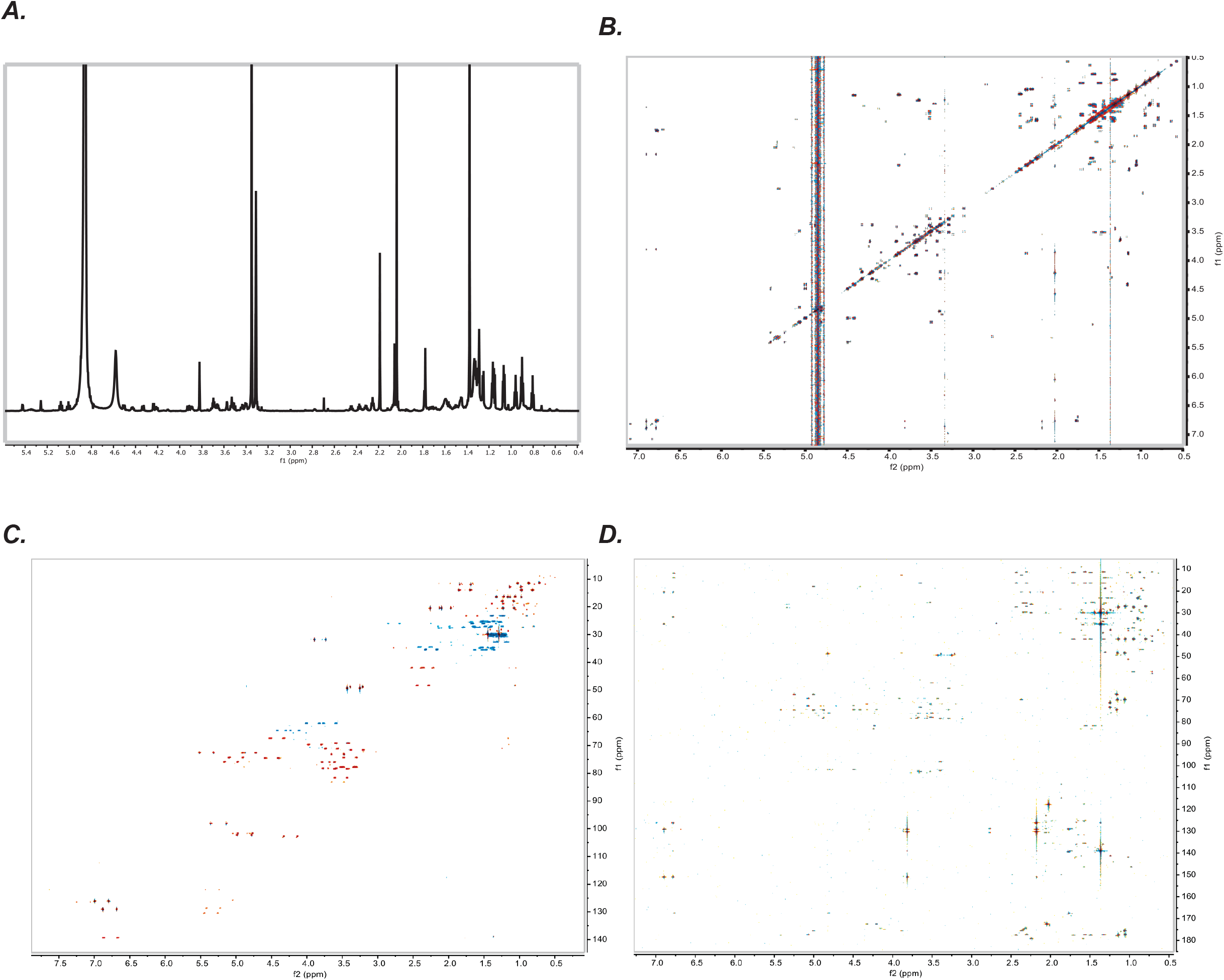
NMR spectra of Dichondrin D (800 MHz, CD_3_OD) (A) ^1^H NMR (B) dqfCOSY (C) HSQC (D) HMBC

**Supplementary Figure 6:**
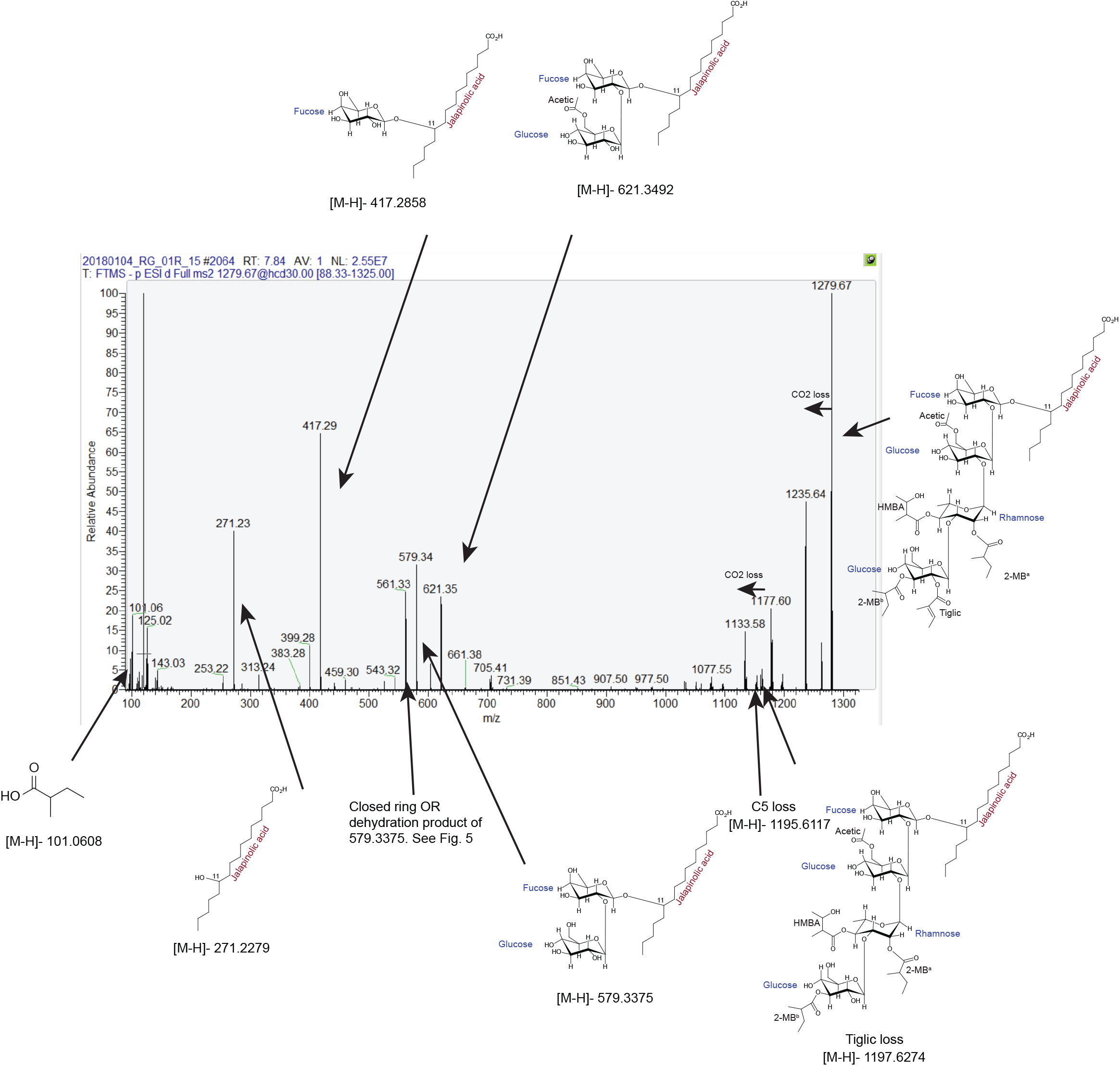
Fragmentation pattern of Dichondrin D. Structures corresponding to various observed MS/MS fragments in negative ion mode are shown. 561.33 is a dehydration/condensation product of the 579.34 fragment. HMBA: Hydroxymethylbutyric acid, 2-MB: 2-methylbutyric acid.

**Supplementary Figure 7:**
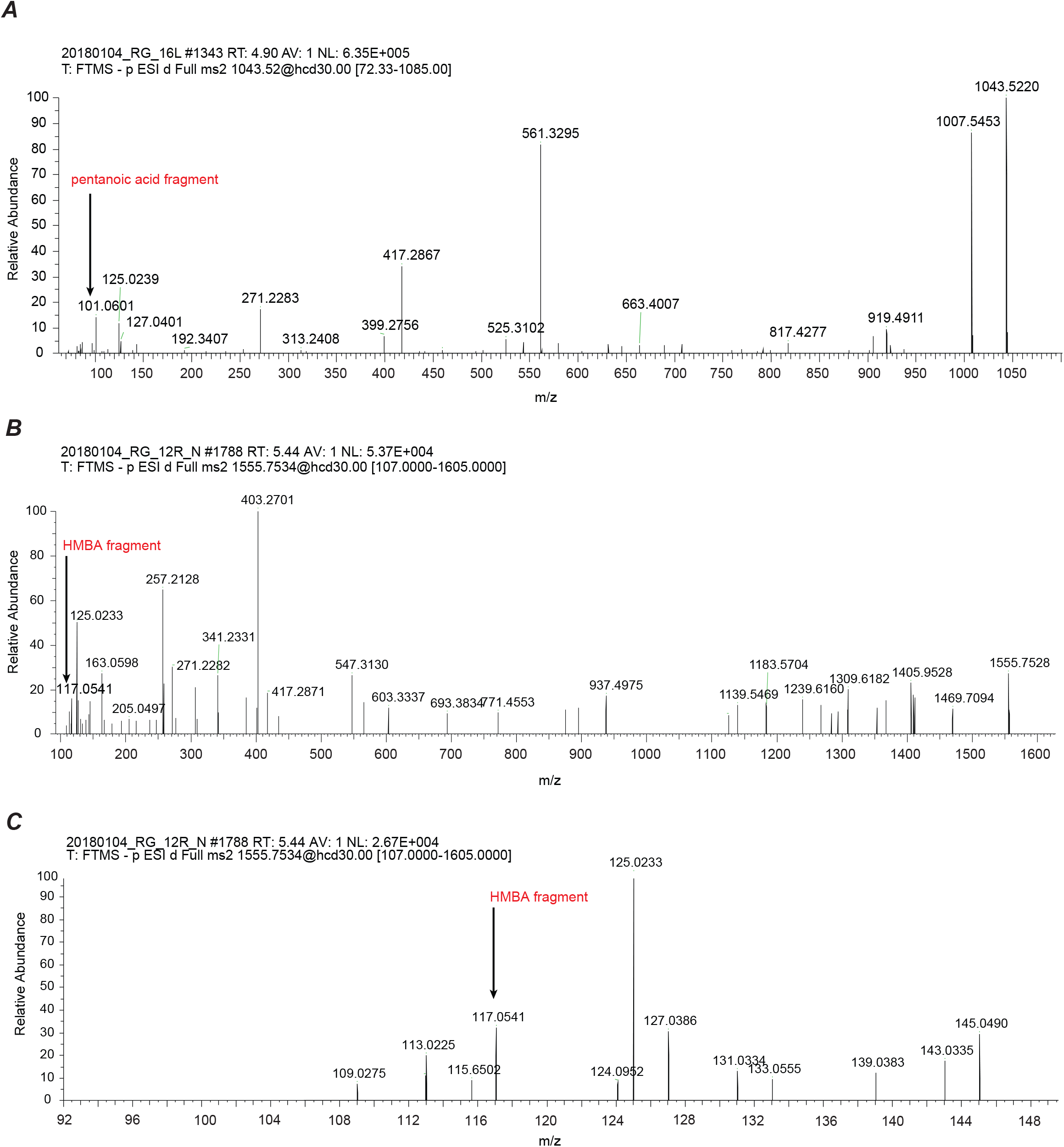
Exemplary MS/MS fragments showing the pentanoic acid and hydroxymethylbutyric acid (HMBA) moiety in *Ipomoea* and *Convolvulus* species. Arrows indicate the respective m/z signal of pentanoic acid and HMBA, respectively. (A) RG (m/z 1043.5220) from *Ipomoea tricolor*. (B) RG from *Convolvulus arvensis* Huntley. (C) A zoomed in view of sub-figure B, showing mass range 92 - 150 m/z.

**Supplementary Fig. 8:**
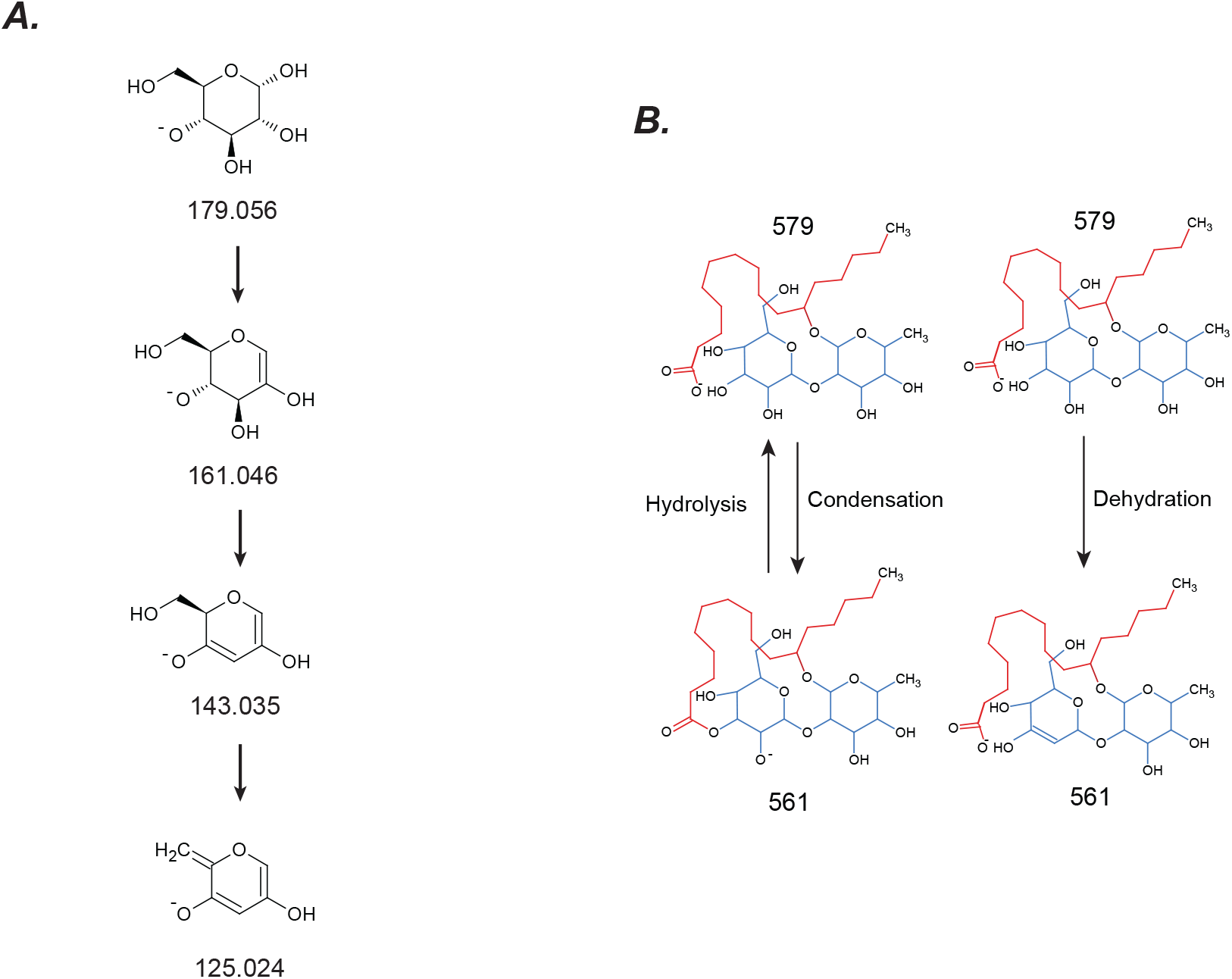
Explaining observed MS/MS fragmentation patterns. All masses denoted are [M-H]-monoisotopic masses of the shown structures (A) Fragmentation pattern of Tricolorin A. (B)Hexose fragmentation producing a 125.024 mass fragment (B) Putative explanations for loss of 18 observed for many neutral losses.

